# Sp1 mechanotransduction regulates breast cancer cell invasion in response to multiple tumor-mimicking extracellular matrix cues

**DOI:** 10.1101/2025.03.18.643983

**Authors:** Abhishek Sharma, Rowan F. Steger, Jen M. Li, Jane A. Baude, Kellie A. Heom, Siddharth S. Dey, Ryan S. Stowers

## Abstract

Breast cancer progression is marked by extracellular matrix (ECM) remodeling, including increased stiffness, faster stress relaxation, and elevated collagen levels. In vitro experiments have revealed a role for each of these factors to individually promote malignant behavior, but their combined effects remain unclear. To address this, we developed alginate-collagen hydrogels with independently tunable stiffness, stress relaxation, and collagen density. We show that these combined tumor-mimicking ECM cues reinforced invasive morphologies and promoted spheroid invasion in breast cancer and mammary epithelial cells. High stiffness and low collagen density in slow-relaxing matrices led to the greatest cell migration speed and displacement. RNA-seq revealed Sp1 target gene enrichment in response to both individual and combined ECM cues, with a greater enrichment observed under multiple cues. Notably, high expression of Sp1 target genes upregulated by fast stress relaxation correlated with poor patient survival. Mechanistically, we found that phosphorylated-Sp1 (T453) was increasingly located in the nucleus in stiff and/or fast relaxing matrices, which was regulated by PI3K and ERK1/2 signaling, as well as actomyosin contractility. This study emphasizes how multiple ECM cues in complex microenvironments reinforce malignant traits and supports an emerging role for Sp1 as a mechanoresponsive transcription factor.

## Introduction

The microenvironment of breast tumors is highly heterogeneous, exhibiting significant spatiotemporal variations in the physical and biochemical properties of the extracellular matrix (ECM)^1^. The progression of breast cancer involves increased collagen deposition, particularly fibrillar collagen I, and enhanced collagen fiber alignment within ECM ^2–4^ (Fig. 1A), resulting in a tumor microenvironment that is significantly stiffer than the healthy breast tissue. The increased collagen density and stiffness serve as an important diagnostic and prognostic marker of disease ^3,5,6^ through manual palpation and radiographic imaging^7^.

**Fig. 1:**
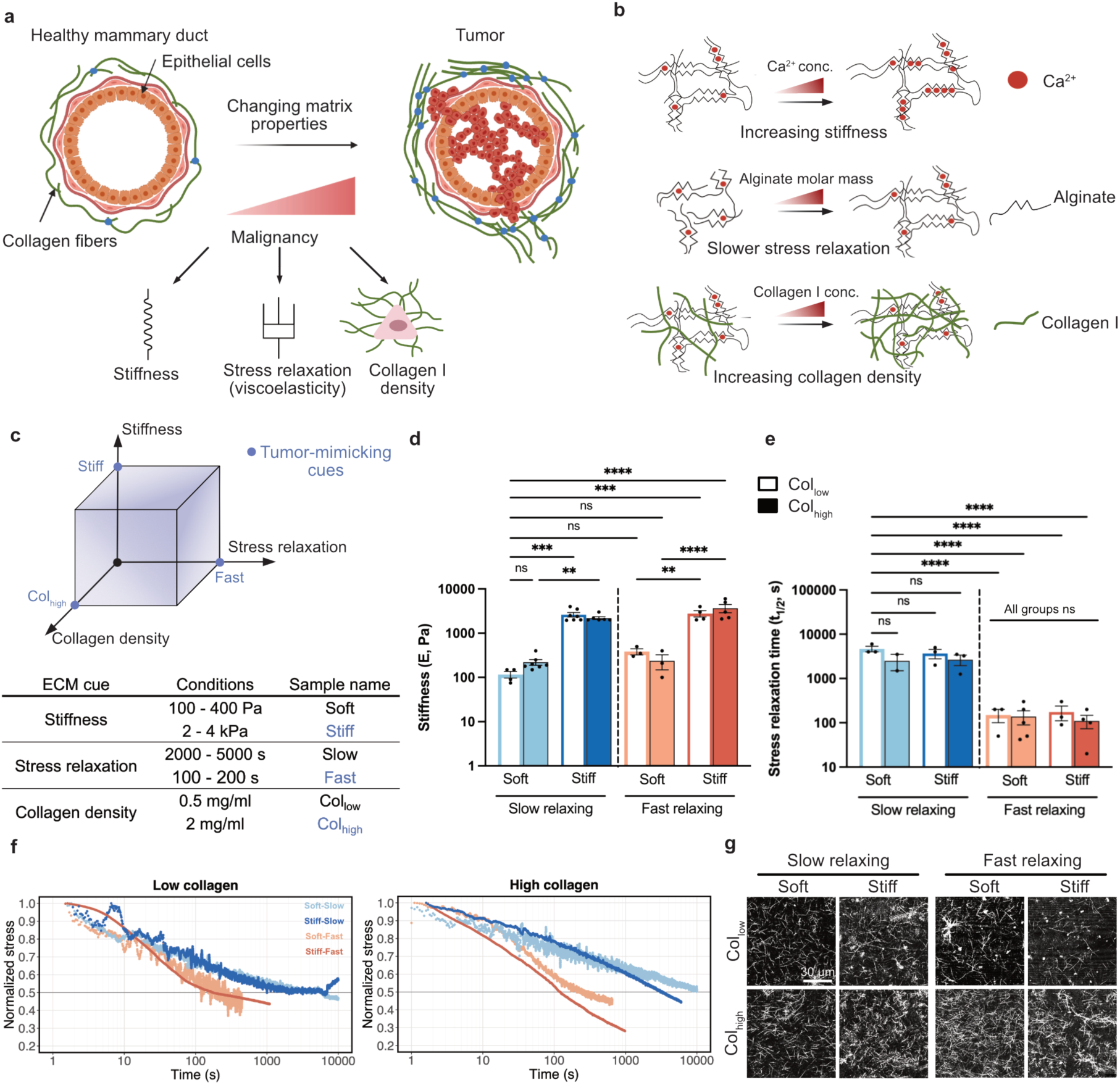
Independent tuning of stiffness, stress relaxation, and collagen fiber density with alginate-collagen I matrices. **a**, Breast cancer progression is accompanied by changes in ECM properties such as elevated stiffness, faster stress relaxation rates, and increased production of collagen I density. **b**, Alginate-collagen matrices allow for independently tuning their stiffness and stress relaxation by varying the calcium ion concentration and the molar mass of the alginate chains, respectively. Further, varying amounts of collagen can be incorporated in these matrices. **c**, We developed 8 matrix conditions as combinations of soft or stiff; slow-relaxing or fast-relaxing, and low or high collagen density. The table describes the nomenclature used throughout the study. **d**, Stiffness measurement of soft and stiff alginate-collagen matrices. (n ≥ 3; mean ± s.e.m.; ANOVA with Tukey’s multiple comparison test). e, Measurement of the time taken to relax stress to half of the maximum value under constant 10% strain (n ≥ 3; mean ± s.e.m.; ANOVA with Tukey’s multiple comparison test). **f**, Stress relaxation curves for alginate-collagen matrices with low and high collagen density. **g**, Confocal reflectance micrographs showing collagen fiber networks within the alginate gel. *p < 0.05; **p < 0.01; ***p < 0.001; ****p < 0.0001, not significant = ns.

Additionally, greater collagen fiber alignment is correlated with poor patient survival rates^8^. Growing evidence also suggests that breast tumors are viscoelastic materials, meaning they relax stress in response to applied deformation^9,10^. Furthermore, differences in viscoelasticity have been shown to discriminate between malignant and benign breast tumors^11,12^. While in vivo studies have documented simultaneous changes in multiple mechanical and biochemical ECM properties, how cells integrate these diverse cues to drive malignant traits remains unclear.

In vitro models have been widely employed to investigate the influence of individual tumor microenvironmental cues on cellular processes including cell cycle progression^9^, invasion^13^, differentiation^14^, and metabolism^15^. Hydrogel-based platforms designed to mimic tumor stiffness have been shown to promote invasion in both breast cancer cells and non-malignant breast epithelial cells^6,16^. Similarly, matrices with faster stress relaxation have been found to induce invasive cell morphologies, enhance migration, and activate cancer-associated signaling pathways in breast epithelial cells^10,17–20^. Additionally, high collagen density and fiber alignment have also been shown to modulate invasion and migration in both normal and malignant cell lines^21–24^. Cells sense these ECM mechanical cues through integrin-mediated focal adhesions and mechanosensitive ion-channels, which transduce signals via the actomyosin cytoskeleton to the nucleus, ultimately driving changes in gene expression^25–27^. Gene expression is regulated, in part, by transcriptional regulators, such as YAP/TAZ and MRTF, that are mechanically responsive^18,28,29^. However, compared to ECM stiffness, our knowledge about how cells sense and respond to ECM viscoelasticity remains severely lacking. Further, while it is well established that individual tumor-mimicking ECM cues can enhance malignant traits in cancer cells, how cells generate an integrated response to multiple cues in complex microenvironments remains poorly understood, primarily because decoupling these cues remains a challenge.

To address this challenge, we utilized alginate-collagen hydrogels with independently tunable stiffness, stress relaxation, and collagen density. We encapsulated mammary epithelial and breast cancer cells, breast cancer spheroids and tumor organoids in these 3D matrices to study the effect of multiple tumor-mimicking ECM cues on cell invasion, migration, and gene expression. We reveal a novel signaling axis for the transcription factor Sp1 that regulates invasion in response to multiple tumor-mimicking cues.

## Results

We aimed to create a platform to tune ECM stiffness, stress relaxation, and collagen I density independently in order to determine how mammary epithelial and breast cancer cells respond when each property is varied alone or collectively. Using an alginate-collagen I interpenetrating network hydrogel, we developed two conditions for each property to mimic healthy or diseased extracellular matrices (Fig. 1B,C): soft or stiff (100 Pa vs. 2-4 kPa)^2^, slow or fast stress relaxing (≈100 s vs. ≈1000 s stress-relaxation half-time)^30,31^, and low or high collagen density (0.5 mg/ml vs. 2 mg/ml)^32–34^. The elastic modulus was varied in these matrices by changing the calcium ion concentration used to ionically crosslink the alginate chains. Stress relaxation time was varied by using alginates of different molecular weights^35^. Collagen density was controlled by incorporating varying amounts of type I collagen into the alginate matrices. Using shear rheology, we confirmed that stiffness could be tuned independent of stress relaxation half-time and collagen density (Fig. 1D). Similarly, stress relaxation half-times could be varied from thousands of seconds in slow relaxing matrices to hundreds of seconds in fast relaxing matrices, independent of the stiffness and collagen density (Fig. 1E,F). Further, upon varying collagen density in these matrices, we saw no significant differences in collagen fiber architecture (Fig. 1G, Supp. Fig. 1A,B). Thus, by varying these three properties over two distinct values, we generated eight unique matrix conditions that can be represented as vertices on a 3D state-space of the tumor microenvironment (Fig. 1C).

Morphology at the single-cell level has been previously shown to correlate with gene expression patterns and tumorigenicity^36–38^. To this end, metastatic breast adenocarcinoma cells (MDA-MB-231) were encapsulated as single cells within these eight matrix conditions and morphologically characterized after 7 days of culture. We measured roundness as the inverse of aspect ratio for individual cells or cell clusters if cells were in contact with neighboring cells. Cells in the Soft-Slow-Col_low_ matrix condition showed highly rounded morphologies and the greatest roundness value among all matrix conditions (Fig. 2A,B).

**Fig. 2:**
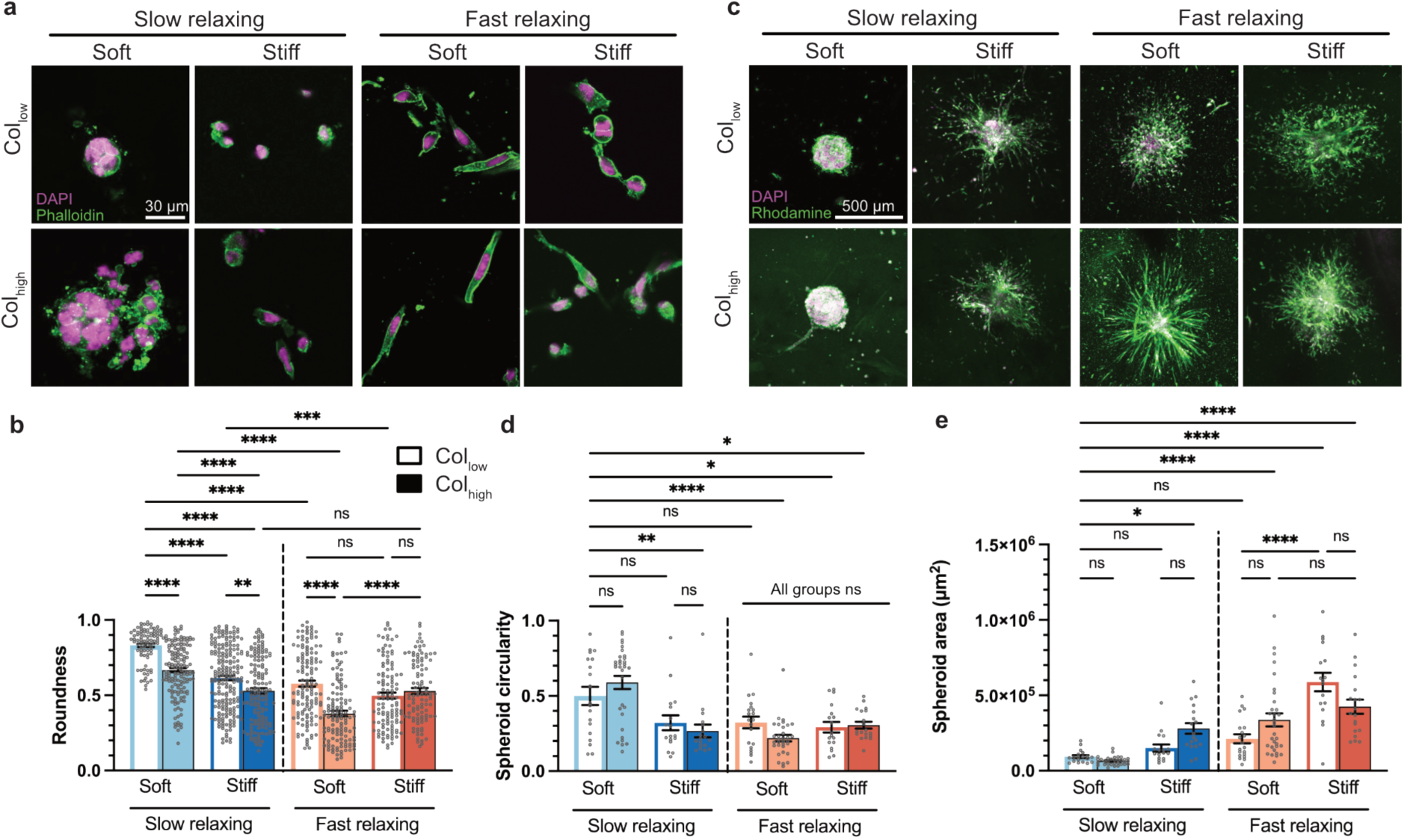
MDA-MB-231 cells and spheroids show invasive morphologies in the presence of individual and multiple tumor-mimicking ECM cues. **a**, Single cells were encapsulated in all 8 alginate-collagen matrix conditions and allowed to grow into clusters over a period of 7 days. **b**, Quantification of cell cluster roundness showed the Soft-Slow-Col_low_ condition led to clusters with the highest roundness. High stiffness, faster stress-relaxation, high collagen or combinations of these cues led to a significant decrease in roundness (n ≥ 20 cells per replicate, with 3 independent replicates; mean ± s.e.m.; Welch ANOVA test with Dunnett’s multiple comparison test). **c**, Spheroids stained with DAPI and Octadecyl Rhodamine-B dye (R-18) were encapsulated and allowed to invade the matrix over a 3-day period. **d, e**, Quantification of spheroid circularity and area. The highest circularity and smallest area were observed in the Soft-Slow-Col_low_ and Soft-Slow-<u>Col</u>_high_ conditions. Compared to Soft-Slow-Col_low_, the presence of two or all three out of high stiffness, faster relaxation, or high collagen led to a significant decrease in circularity and increase in area. (n ≥ 5 spheroids per replicate, with 3 independent replicates; mean ± s.e.m., ANOVA with Tukey’s multiple comparison test). *p < 0.05; **p < 0.01; ***p < 0.001; ****p < 0.0001, not significant = ns.

Compared to the Soft-Slow-Col_low_ condition, the presence of high stiffness, fast stress relaxation, or high collagen density individually, led to a statistically significant reduction in cell or cluster roundness (≈26%, 31%, and 20%, respectively). This is in agreement with previous studies showing that epithelial cells adopted invasive morphologies in response to either of these cues^6,18,30,39,40^. We observed even larger reductions in cluster roundness with respect to the Soft-Slow-Col_low_ condition when two or all three cues were presented collectively at tumor-mimicking levels compared to just one. For example, the presence of both high stiffness and fast relaxation led to ≈ 40% decrease in roundness. Thus, the combined effect of multiple tumor-mimicking ECM cues has a greater impact on cell morphology than any one individual cue, demonstrating a synergistic interaction between ECM cues. Additionally, we observed instances where a tumor-mimicking cue did not significantly affect cluster roundness if another cue was already present. For instance, in Stiff-Fast-Col_low_ versus Stiff-Fast-Col_high_ matrices, the presence of high collagen density showed no significant decrease in roundness. Similarly, in Stiff-Slow-Col_high_ matrix versus Stiff-Fast-Col_high_ matrices, fast stress relaxation also led to no significant change in cluster roundness. These findings suggest that, in certain contexts, a single tumor-mimicking cue may not be sufficient to override the influence of other ECM cues in driving changes in cell morphology.

We also repeated this study in a tumorigenic but non-metastatic cell line (MCF-7) and a non-tumorigenic cell line (MCF-10A) to understand whether other mammary epithelial cell lines of different tumorigenicity show similar changes in cell morphology in response to multiple tumor-mimicking cues. In both these cell lines, we also observed a significant decrease in cell cluster roundness in response to one or multiple cues compared to the Soft-Slow-Col_low_ condition (Supp. Fig.2A-D). Out of all three cues, MCF-10A cells showed the greatest decrease in roundness in response to high collagen (≈27%) in comparison to fast relaxation (≈17%) or high stiffness (≈14%). In contrast, MDA-MB-231 cells showed the most pronounced reduction in roundness in response to fast stress relaxation. Furthermore, MCF-10A cells showed a distinct sensitivity to collagen concentration, with significant differences in roundness between the Stiff-Fast-Col_low_ and Stiff-Fast-Col_high_ groups, an effect not observed in MDA-MB-231 cells. These findings suggest that different cell lines exhibit varying sensitivities towards tumor-mimicking ECM cues.

To investigate whether changes in morphology from encapsulated single cells were indicative of an invasive phenotype, we encapsulated MDA-MB-231 spheroids in alginate-collagen matrices and analyzed their invasiveness after 3 days of culture. Spheroid invasion was evaluated using circularity, calculated as the ratio of spheroid area to the square of its perimeter, which captures shape irregularities during invasion. We saw that MDA-MB-231 spheroids in the Soft-Slow-Col_low_ and Soft-Slow-Col_high_ matrix conditions were not invasive and showed the highest circularity and smallest invasion area (Fig. 2C,D). Notably, compared to the Soft-Slow-Col_low_ condition, the presence of high stiffness, fast relaxation, or high collagen density individually led to no significant changes in spheroid circularity or area. (Fig. 2D,E). The presence of two or all three tumor-mimicking cues, however, led to a significant decrease in circularity and increase in spheroid area (Fig. 2C,D,E). Thus, the presence of multiple tumor-mimicking ECM cues has a greater impact on the cell invasive phenotype than individual cues.

MCF-10A spheroids also showed a significant decrease in circularity and increase in area in response to multiple cues (Supp. Fig. 2E,F,G). MCF-10A spheroids in Soft-Slow-Col_low_ matrices showed a significantly larger decrease in circularity in the presence of high collagen density (≈30%) compared to MDA-MB-231 spheroids (≈2%) (Fig. 2D, Supp. Fig. 2E,F). This highlights their greater sensitivity towards high collagen density, similar to the trends seen in single cell encapsulation studies (Supp. Fig. 2D).

We also encapsulated murine mammary organoids in all 8 matrix conditions and observed changes in their morphology over a period of 2 weeks. Consistent with our single cell and spheroid encapsulation studies, we observed that the organoids remained round and did not invade the matrix in the Soft-Slow-Col_low_ condition (Supp. Fig.2H). While organoids were less rounded and showed an invasive morphology in all stiff conditions, in the Soft-Slow-Col_high_ condition, the organoids elongated and branched similar to events observed during mammary morphogenesis in organoid cultures^41,42^. This suggests that mammary organoids can undergo invasion or morphogenesis depending on the presence of specific ECM cues and their combinations in their environment.

Increased collagen fiber alignment is a prognostic signature of poor survival in breast cancer patients^8^. We therefore determined if cells also were also aligning the collagen fibers during invasion. Collagen fiber alignment was measured with respect to the cell boundary in all 8 matrix conditions after 7 days of culture. We quantified aligned collagen fibers as the proportion of total fibers oriented at an angle greater than 70° relative to the cell boundary.

We observed that MDA-MB-231 cells showed the smallest fraction of aligned fibers in the Soft-Slow-Col_low_ condition (Supp. Fig. 3A,B). All other matrix conditions showed a significantly higher fraction of aligned fibers. MCF-10A cells also showed a significantly higher percentage of aligned fibers compared to the Soft-Slow-Col_low_ group, except in the Stiff-Slow-Col_high_ condition (Supp. Fig. 3C). This observed reduction in fiber alignment in our high collagen matrix condition is in line with previous studies showing decreased collagen fiber orientation and cell alignment in high collagen matrices^43,44^.

We next investigated the extent to which stiffness, stress relaxation, and collagen density impact migration of MDA-MB-231 metastatic breast cancer cells. Using time-lapse confocal microscopy, we tracked cell positions over 16 hours and found that cells in Soft-Slow-Col_low_ and Soft-Slow-Col_high_ conditions showed the lowest mean squared displacement (MSD) and average speed. Compared to the Soft-Slow-Col_low_ matrix condition, the presence of high stiffness led to the highest increase in MSD and average speed (Fig. 3A-C). While fast stress relaxation led to a significant increase in average speed but not MSD, high collagen led to no significant changes in neither MSD nor average speed. Cells in Soft-Slow-Col_low_ and Soft-Slow-Col_high_ conditions showed rotation around their centroid but no translation (Supp. Video 1,5), in line with previous studies^45,46^. Notably, faster stress relaxation promoted cellular protrusions that dynamically extended and retracted over time (Fig. 3A, Supp. Video 3), as previously reported^10,17^. In many cases, these dynamic protrusions did not lead to persistent migration, as measured by translation of the centroid of the cell and thus are not accounted for in our MSD and speed metrics. Further, in the collective presence of high stiffness and fast stress relaxation, cells showed both increased stiffness-driven enhanced migration (Fig. 3A), as well as fast relaxation-driven formation of dynamic protrusions (Supp. Video 4,8). In Stiff-Slow-Col_low_ versus Stiff-Fast-Col_low_ conditions, the presence of fast stress relaxation induced a significant reduction in both MSD and speed (≈70% MSD and ≈87% average speed) (Fig. 3B,C,D). Similarly, compared to the Stiff-Slow-Col_low_ condition, the presence of high collagen density also led to a large significant decrease in both MSD (≈936%) and speed (≈107%). This was also true in fast relaxing matrices, where, compared to the Stiff-Fast-Col_low_ condition, the presence of high collagen density also led to a significant decrease (≈57%) in average speed and a decrease in MSD (≈184%), though that change was not statistically significant. This observed reduction in cell migration in response to high collagen density is in line with previous studies^47^. In summary, we observe that the presence of multiple tumor-mimicking cues can lead to cell populations that display diverse phenotypic behaviors. Our results also depict that in complex microenvironments, cell migration in response to a specific ECM cue can be strongly influenced by the concurrent presence of other ECM cues.

**Fig. 3:**
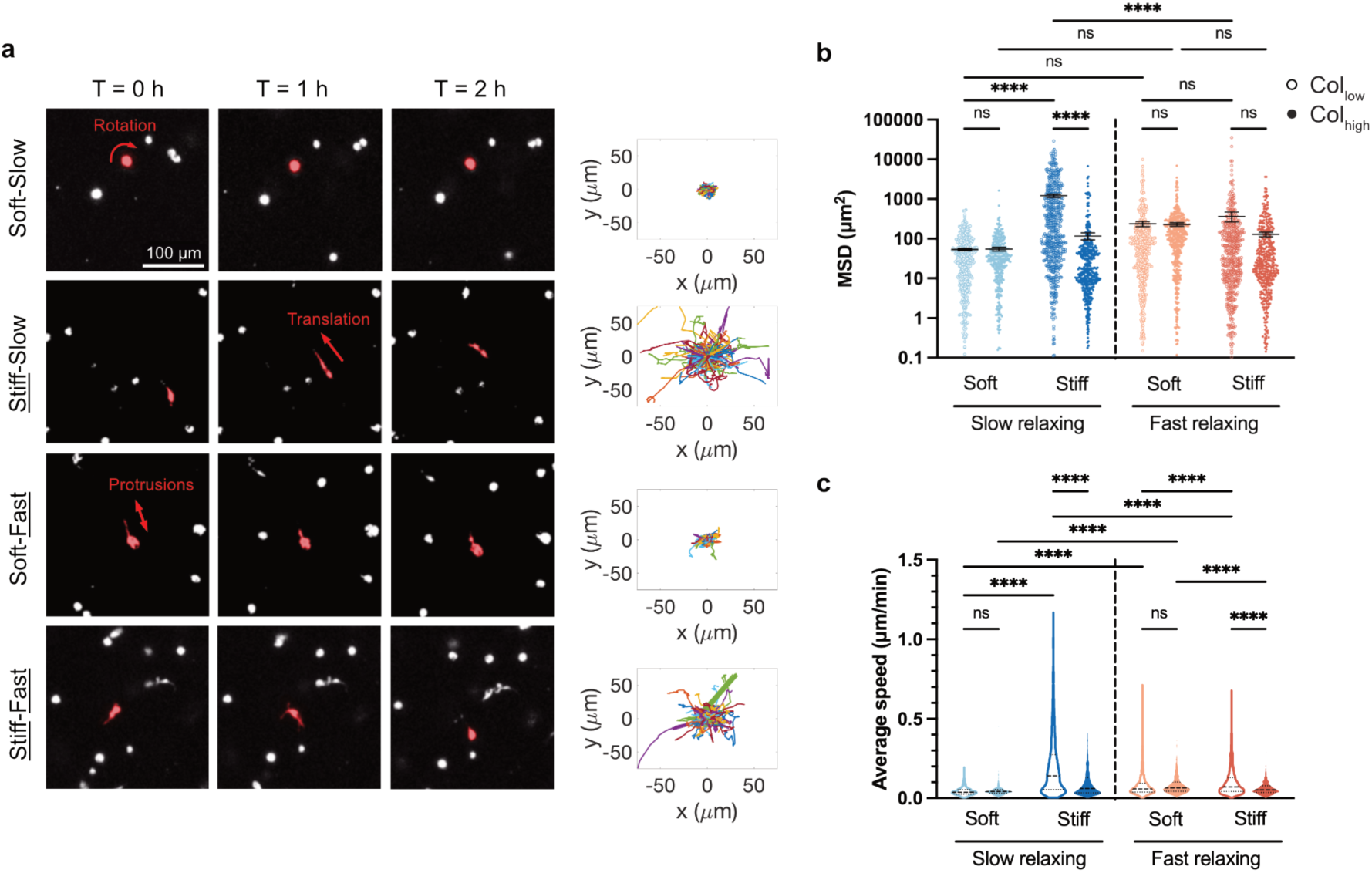
Matrix stiffness drives the largest increases in cell migration compared to other tumor-mimicking ECM cues. **a**, Live-cell imaging of MDA-MB-231 cells in Col_low_ matrices that were Soft-Slow, <u>Stiff</u>-Slow, Soft-<u>Fast</u>, or <u>Stiff</u>-<u>Fast</u>. The centroid of cells was tracked and their migration trajectory projection in the X-Y plane is plotted. **b**, Mean squared displacement (MSD) and **c**, average speed were measured across all 8 matrix conditions. The greatest increase in MSD and average speed was observed in the <u>Stiff</u>-Slow-Col_low_ condition. n ≥ 100 cells, with 3 independent replicates; mean ± s.e.m.; ANOVA with Tukey’s multiple comparison test. *p < 0.05; **p < 0.01; ***p < 0.001; ****p < 0.0001, not significant = ns.

To investigate how gene expression is influenced by stiffness, stress relaxation, and collagen density, we performed RNA sequencing on cells cultured for 7 days in all eight matrix conditions. Differential gene expression analysis was conducted by comparing each condition to the Soft-Slow-Col_low_ baseline. In a scenario where each ECM cue independently regulates a distinct subset of genes, one would expect that the number of differentially expressed genes in the presence of multiple ECM cues would be equal to the sum of the differentially expressed genes from individual cues. Interestingly, with MDA-MB-231 cells, we found many cases where the presence of multiple ECM cues led to many more differentially expressed genes than the sum of those from the individual cues (Fig. 4A). For example, Stiff-Slow-Col_low_ matrix (high stiffness) led to 239 differentially expressed genes and Soft-Slow-Col_high_ (high collagen) led to 63 differentially expressed genes. However, when both high stiffness and high collagen were collectively present in the Stiff-Slow-Col_high_ matrix, we observed 895 differentially expressed genes, many of which overlapped with those from the Stiff-Slow-Col_low_ and Soft-Slow-Col_high_ conditions (Fig. 4A–C). In contrast, for MCF-10A cells, the number of differentially expressed genes enriched in the presence of multiple tumor-mimicking cues was not higher than the sum of differentially expressed genes enriched in response to the individual cues, thus suggesting a lack of transcriptional synergy in these cells (Supp. Fig. 4A). The greater effect on gene expression in the presence of multiple cues in metastatic MDA-MB-231 cells might be due to their higher genomic instability^48,49^, signaling pathway crosstalk^50^, as well as a greater genetic and phenotypic heterogeneity^51,52^.

**Fig. 4:**
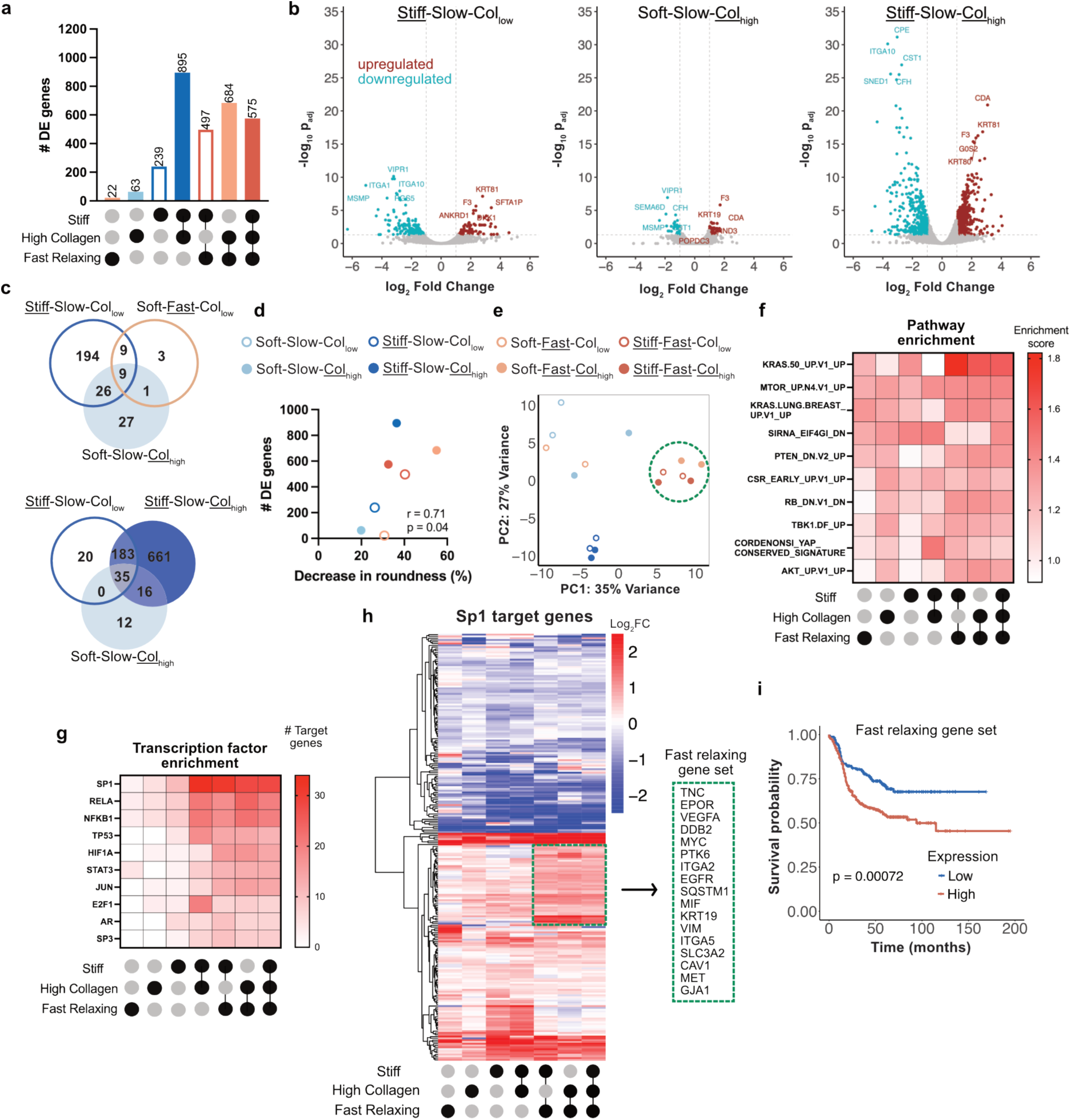
Tumor mimicking cues are associated with enrichment of Sp1 target genes in MDA-MB-231 cells. **a**, The number of differentially expressed genes with respect to the Soft-Slow-Col_low_ condition was determined for all matrix conditions, revealing that multiple ECM cues induce a greater number of differentially expressed genes than the sum of individual cues (n = 2 replicates). **b**, Volcano plot of gene expression for <u>Stiff</u>-Slow-<u>Col</u>_high_ condition shows a greater log_2_ fold change and – log_10_p_adj_ than in <u>Stiff</u>-Slow-Col_low_ and Soft-Slow-<u>Col</u>_high_ conditions. **c**, Venn diagram plots show little overlap in gene enrichment between high stiffness, fast relaxation, and high collagen conditions. Stiff-Col_high_ conditions showed a large overlap in gene enrichment with both Stiff-Col_low_ and Soft-Col_high_ conditions. **d**, Mapping the number of differentially expressed genes for a given matrix condition against percentage decrease in cell cluster roundness shows a positive correlation. One-tailed Spearman correlation test was used to determine significance. **e**, PCA analysis shows that all fast relaxing groups (except Soft-<u>Fast</u>-Col_low_) clustered together. **f**, Heat map for GSEA Onco-DB pathway enrichment analysis for each matrix condition shows enrichment of KRAS signaling. **g**, Heat map showing top transcription factors enriched using TRRUST analysis for each matrix condition. Sp1 showed enrichment in all matrix conditions and had the largest number of target gene enrichment. **h**, Mapping Sp1 target gene expression identified 17 genes in the fast relaxing cluster overlapping with known Sp1 targets associated with malignant neoplasm and metastasis of breast. **i**, Recurrence-free survival analysis for n = 422 patients with basal PAM50 subtype from the TCGA using the upregulated fast relaxing gene set showed a significantly decreased survival probability. p-value was calculated using the logrank test.

Notably, when analyzing the effects of individual ECM cues, the differentially expressed gene subsets from individual high stiffness, fast relaxation, and high collagen conditions were largely mutually exclusive in MDA-MB-231 cells, with only a small fraction overlapping between these matrices (Figure 4C). Further, we also observed a positive correlation between the number of differentially expressed genes in response to single or multiple tumor-mimicking cues and the extent of reduction in cell cluster roundness in both MDA-MB-231 and MCF-10A cells (Fig. 4D, Supp. Fig.4B). Performing principal components analysis (PCA) on all 8 matrix conditions revealed that with MDA-MB-231 cells, fast relaxing matrix conditions (except Soft-Fast-Col_low_) clustered together (Fig. 4E). However, with MCF-10A cells, all high collagen conditions were clustered together, consistent with their greater collagen sensitivity (Supp. Fig. 4C). In summary, our findings demonstrate that the integration of multiple tumor-mimicking cues by cells can elicit a broader transcriptional response than the sum of their individual effects. Furthermore, the magnitude of these gene expression changes correlates with the extent of downstream alterations in invasive cell morphology.

We next performed Gene Set Enrichment Analysis (GSEA) against the oncogenic signature gene set (C6) in the Human Molecular Signatures Database (MSigDB)^53,54^. Our analysis revealed that KRAS signaling, associated with disease progression in multiple solid cancers^55^, was enriched in both MDA-MB-231 and MCF-10A cells in all conditions marked by high stiffness, fast relaxation, or high collagen, as well as combinations of these cues (Fig. 4F, Supp. Fig. 4D). This shows that our alginate-collagen matrices with tunable ECM cue parameters lead to enrichment of multiple signaling pathways implicated in breast cancer.

We investigated transcription factor enrichment across various matrix conditions relative to the Soft-Slow-Col_low_ condition using the TRRUST database^56^. Several well-known transcription factors associated with breast cancer were enriched (Fig. 4G, Supp. Fig. 4E). Among these, we found that Sp1, whose expression is inversely correlated with patient survival in multiple cancers^57^, showed a high enrichment of target genes in both MDA-MB-231 and MCF-10A cells in response to high stiffness, fast relaxation, high collagen density, as well as the combination of these cues. Additionally, the number of enriched Sp1 target genes was higher in matrix conditions incorporating multiple tumor-mimicking cues compared to those with individual cues. This strong enrichment of Sp1 target genes aligns with prior studies demonstrating its critical role in mediating cellular response to stiff matrices and direct mechanical force application, further emphasizing its importance in mechanoresponsive pathways^16,58^.

Notably, hierarchical clustering of Sp1 target genes in MDA-MB-231 cells revealed a gene set that was also upregulated in all fast relaxing conditions except for the Soft-Fast-Col_low_ condition (Fig. 4H). On comparing this upregulated gene cluster to known Sp1 target genes involved in breast neoplasm and neoplasm metastasis, we saw an overlap of 17 target genes. Using mRNA expression data from The Cancer Genome Atlas (TCGA)^59^ for patients with the PAM50 basal subtype, we observed that patients with a higher expression of these genes exhibited a significantly reduced recurrence-free survival probability (Fig. 4I). Patients with a high expression had a mean survival time of 18 months, while for patients with a low gene expression, this time was 45.4 months. Taken together, our results suggest that upregulation of Sp1 target genes in fast stress relaxing conditions is associated with poor patient survival.

Since we saw Sp1 target gene enrichment in all matrix conditions that induced invasion, we sought to determine its role in regulating the malignant phenotype. Although Sp1 is known to play a role in cancer progression^57^, its role as a mechanotransducer is understudied. To this end, we used mithramycin-A, a well-established small molecule inhibitor of Sp1^60,61^ to treat MDA-MB-231 and MCF-10A cells encapsulated in matrices with high stiffness, fast relaxation, or the presence of both these mechanical cues (Stiff-Slow, Soft-Fast, and Stiff-Fast). Sp1 inhibition led to a significant increase in cell roundness for both cell types in all three treated groups compared to the DMSO vehicle controls (Fig. 5A,B, Supp. Fig. 5A,B).

**Fig. 5:**
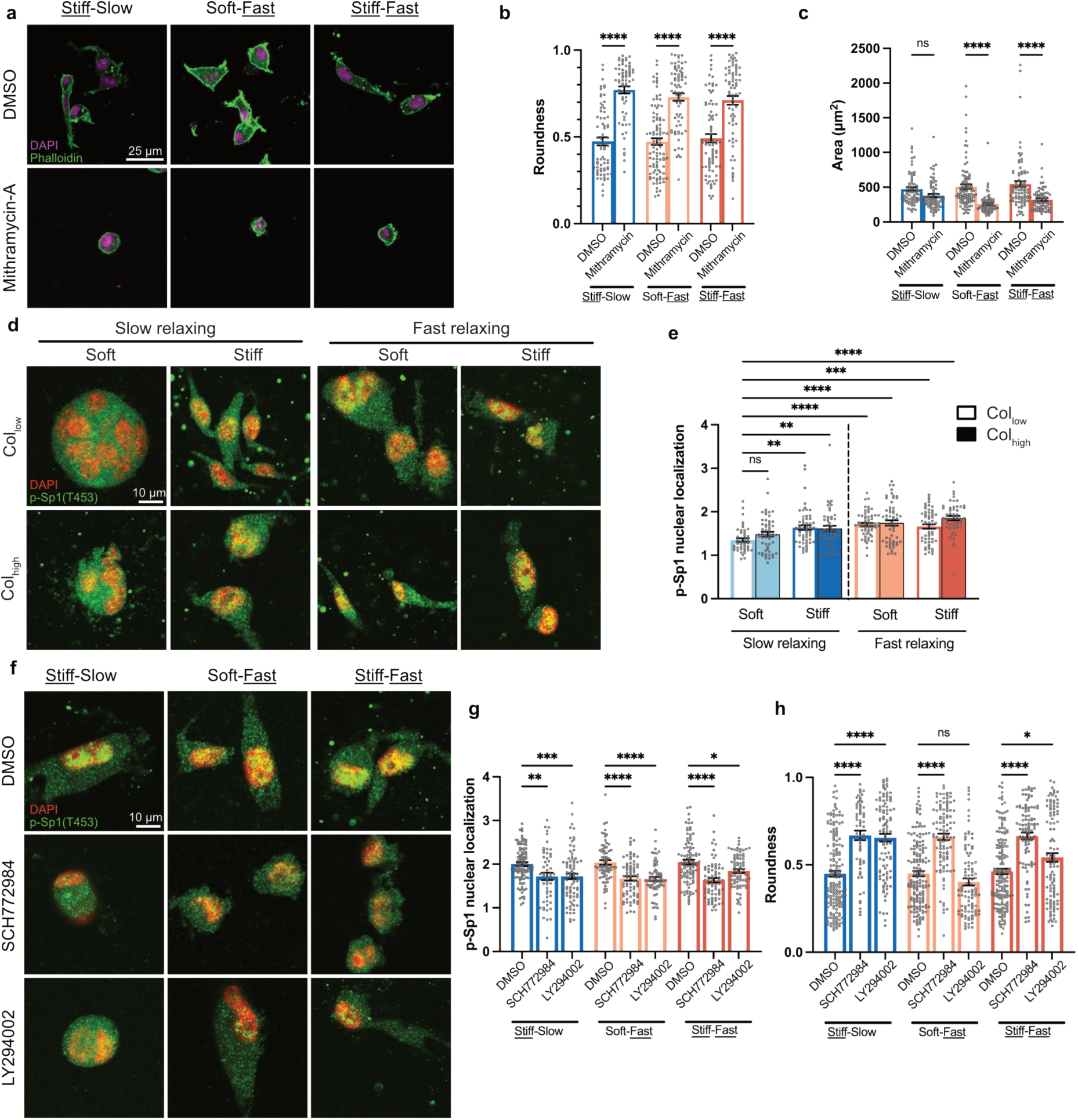
Sp1 induces the invasive phenotype in response to multiple tumor mimicking ECM cues. **a**, Inhibiting Sp1 activity using mithramycin-A led to a significant increase in MDA-MB-231 cluster roundness in <u>Stiff</u>-Slow, Soft-<u>Fast</u>, as well as <u>Stiff</u>-<u>Fast</u> matrix conditions. **b, c** Quantification of the change in roundness and cluster area upon Sp1 inhibition. (n ≥ 15 cells per replicate, with 3 independent replicates; ANOVA with Sidak’s multiple comparison test). **d**, Immunofluorescence imaging of p-Sp1(T453) and **e**, quantification of the nuclear to cytoplasmic intensity levels shows a significant increase in most matrix conditions compared to the Soft-Slow-Col_low_ condition. (n ≥ 15 cells, with 3 independent replicates; mean ± s.e.m; ANOVA with Dunnett’s multiple comparison test). **f**, Confocal immunofluorescence imaging of cells treated with ERK1/2 inhibitor (SCH772984) and PI3K inhibitor (LY294002). **g**, Both ERK1/2 and PI3K inhibition led to a significant decrease in nuclear localization of p-Sp1(T453) in Stiff-Slow, Soft-<u>Fast</u>, as well as <u>Stiff</u>-<u>Fast</u> matrix conditions. **h**, Cluster roundness was quantified for phalloidin stained cells. While ERK1/2 inhibition led to significant decrease in all conditions, PI3K inhibition had a significant effect only in <u>Stiff</u>-(Slow and <u>Fast</u>) matrices. For both **g, h**, n ≥ 15 cells per replicate, with 3 independent replicates; mean ± s.e.m.; ANOVA with Sidak’s multiple comparison test. *p < 0.05; **p < 0.01; ***p < 0.001; ****p < 0.0001, not significant = ns.

While MDA-MB-231 cells showed a significant decrease in cluster area in fast relaxing matrices (Fig. 5C), MCF-10A cells showed a significant decrease in area in all three treatment conditions (Supp. Fig. 5C).

Increased phosphorylation of Sp1 at Thr-453 has been previously associated with formation of invasive cell clusters in stiff alginate-Matrigel hydrogels^16^. Further, Sp1 phosphorylation has been shown to increase its transcriptional activity^62–64^. To this end, we investigated whether there are changes in levels of Sp1 phosphorylation (T453) in response to tumor-mimicking ECM cues that drive the invasive phenotype. Notably, in MDA-MB-231 cells, we observed that compared to the Soft-Slow-Col_low_ condition, there was a significant increase in levels of nuclear-to-cytoplasmic ratio of phospho-Sp1 in all matrix conditions (except Soft-Slow-Col_high_ condition), thus correlating with the occurrence of invasive morphologies (Figure 5D,E). MCF-10A cells, however, showed high levels of phospho-Sp1 nuclear localization even in the Soft-Slow-Col_low_ condition, and no significant changes in response to most ECM cues, except in the Soft-Fast-Col_low_ and Stiff-Slow-Col_high_ conditions (Supp. Fig. 5D,E). We believe that high basal levels of phospho-Sp1 in these cells and their inherently non-malignant phenotype likely contribute to the observed differences in phospho-Sp1 nuclear localization.

Both ERK1/2 and PI3K are known to phosphorylate Sp1 at Thr-453 and in various cell types, including breast cancer cells^65–68^. Interestingly, it has been shown that Sp1 is differentially phosphorylated in response to 3D matrix stiffness, as well as upon direct mechanical force application^16,58^. We therefore asked whether phosphorylation of Sp1 at Thr-453 by ERK1/2 or PI3K is critical for driving invasion in response to tumor-mimicking ECM cues. Both ERK1/2 inhibition with SCH77298 and PI3K inhibition with LY294002 led to a significant decrease in phospho-Sp1 nuclear localization in MDA-MB-231 and MCF-10A cells in response to high stiffness, fast relaxation, or the presence of both these mechanical cues (Fig. 5F,G, Supp. Fig. 5G). Further, ERK1/2 inhibition also led to a significant increase in cluster roundness in all three mechanical conditions in both cell lines (Figure 5H, Supp. Fig. 5H). While PI3K inhibition in MDA-MB-231 cells led to a significant increase in roundness in stiff matrices independent of stress relaxation (Stiff-Slow, Stiff-Fast), MCF-10A cells showed a significant increase in roundness in slow relaxing (Stiff-Slow) condition only. Our results show that Sp1 mechanosignaling drives the malignant phenotype in response to specific tumor-like matrix mechanical properties via phosphorylation at Thr453 by ERK1/2 and PI3K.

The actin cytoskeleton has been previously shown to affect cell response to ECM mechanical cues either via direct force transmission to the nucleus or through downstream signaling cascades^25,69^. We therefore asked whether actomyosin contractility also regulates the occurrence of invasive morphologies in breast cancer cells via Sp1 signaling. Since contractility in non-muscle cells is driven by phosphorylation of myosin via myosin light chain kinase (MLCK)^70–72^, we used a MLCK inhibitor, ML-7, to disrupt cell contractility. In ML-7 treated groups, both MDA-MB-231 and MCF-10A cells showed a significant increase in roundness, but only in the Stiff-Slow matrix condition, with no significant effect observed in fast relaxing matrices (Soft-Fast, Stiff-Fast) (Figure 6A,C, Supp. Fig. 6A,B). Since it has been previously demonstrated that cells respond to fast relaxing environments by forming actin-rich protrusions^10,17,18,73^, we disrupted actin network polymerization using Cytochalasin-D. Compared to DMSO-treated controls, both MDA-MB-231 and MCF-10A cells treated with Cytochalasin-D showed a significant increase in roundness in the presence of high stiffness, as well as fast stress relaxation (Stiff-Slow, Soft-Fast, Stiff-Fast). Notably, MDA-MB-231 cells treated with either ML-7 or Cytochalasin-D showed a significant decrease in nuclear localization of phospho-Sp1 only in stiff matrices (Stiff-Slow and Stiff-Fast), but not in the Soft-Fast matrix (Figure 6 A,D). MCF-10A cells, however, exhibited a unique response where ML-7 treatment led to reduced phospho-Sp1 nuclear localization only in the slow relaxing, stiff matrix condition (Stiff-Slow), however, Cytochalasin-D treatment led to reduced nuclear localization across all three conditions, in line with their increased roundness (Supp. Fig. 6A-C). Thus, actin polymerization regulates phospho-Sp1 nuclear localization in MCF-10A cells across stiff, as well as fast relaxing tumor-mimicking mechanical conditions, whereas in MDA-MB-231 cells, this regulation is specific to stiff environments. In summary, we observed that induction of invasive cell morphologies occurs in a mechanical cue and cell-line-specific manner via myosin contractility and actin polymerization, which drives downstream changes in phospho-Sp1 nuclear localization.

**Fig. 6:**
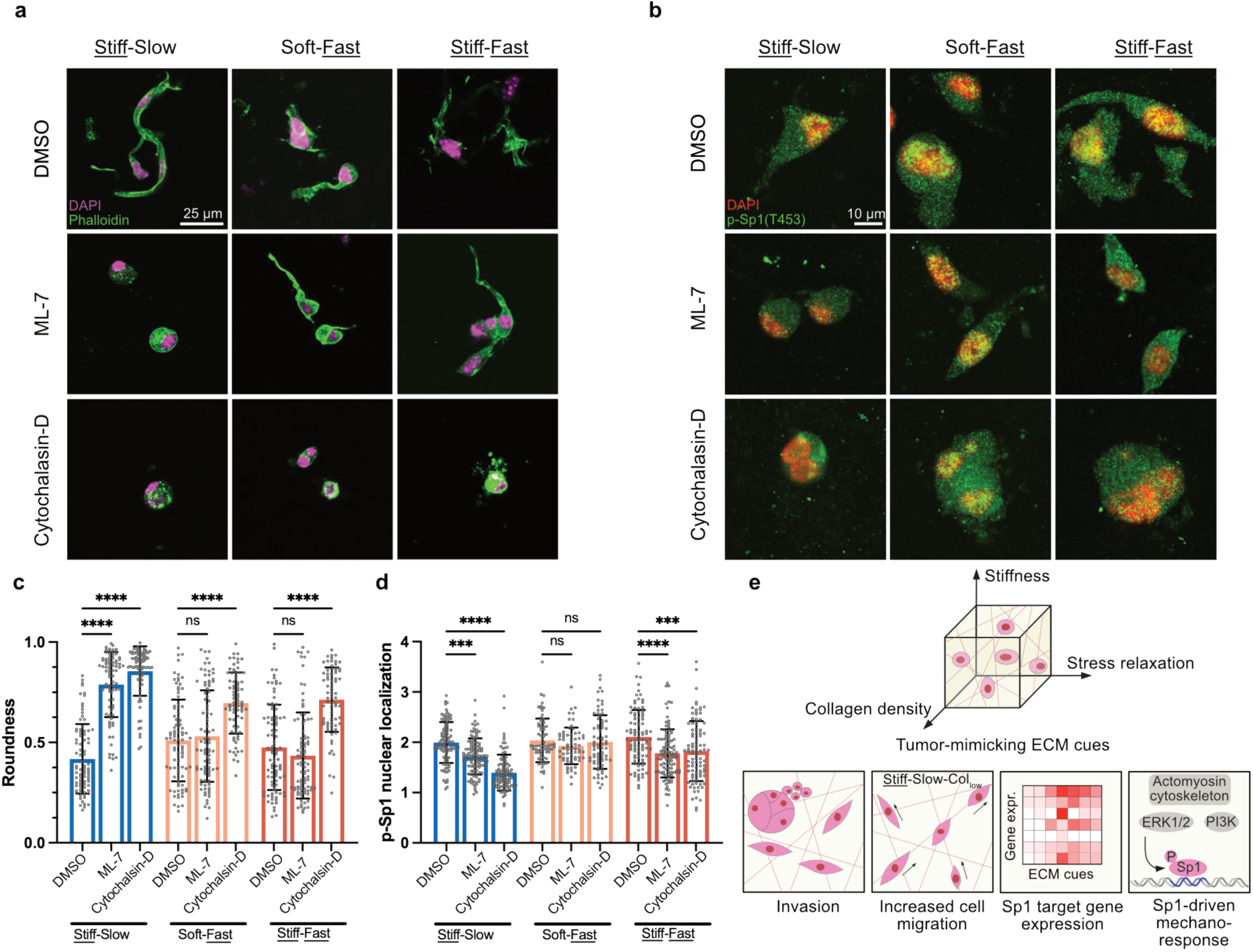
Actomyosin cytoskeleton regulates p-Sp1 nuclear localization in stiff matrices. **a**, **b** Myosin light chain kinase (MLCK) was inhibited via ML-7 and actin polymerization was inhibited via cytochalasin in MDA-MB-231 cells encapsulated in <u>Stiff</u>-Slow, Soft-<u>Fast</u>, as well as <u>Stiff</u>-<u>Fast</u> matrix conditions. Immunofluorescence imaging was performed on phalloidin, DAPI, and p-Sp1(T453) stained MDA-MB231 cells. **c**, Inhibition of MLCK led to a significant increase in cluster roundness only in <u>Stiff</u>-Slow relaxing condition, and inhibition of actin polymerization led to a significant increase in roundness in all 3 conditions. **d**, MLCK and actin polymerization inhibition led to a significant decrease in nuclear localization levels of p-Sp1 only in stiff matrices (<u>Stiff</u>-Slow and <u>Stiff</u>-<u>Fast</u>). **e**, Summary schematic showing that in complex environments with multiple tumor-mimicking ECM cues such as high stiffness, altered stress relaxation, and high collagen density, cells undergo changes in their morphological and migratory phenotype. Multiple ECM cues can also act synergistically at the transcriptional and signaling pathway level. We observed enrichment of Sp1 target genes in response to multiple ECM cues. Sp1 phosphorylation is regulated via the actomyosin machinery and ERK1/2 and PI3K kinases, which drives the downstream malignant traits. For both **c, d**, n ≥ 15 cells per replicate, with 3 independent replicates; mean ± s.e.m.; ANOVA with Sidak’s multiple comparison test. *p < 0.05; **p < 0.01; ***p < 0.001; ****p < 0.0001, not significant = ns. All 3 conditions have a low collagen density.

## Discussion

The tumor ECM presents a heterogeneous microenvironment where a complex interplay of mechanical and biochemical cues collectively governs cellular invasion within the tissue. Our reductionist approach, using alginate-collagen hydrogels with independently tunable properties, has allowed us to systematically vary stiffness, stress relaxation rates, and collagen density, providing a controlled platform to dissect their individual and combined effects. Using multiple cell lines, spheroids, and murine mammary organoids, we show that ECM physical and biochemical cue combinations can generate diverse phenotypes. We identified that a decrease in cell and cluster roundness is reinforced when multiple tumor-associated cues are present in the microenvironment surrounding the cells. In MDA-MB-231 cells, this trend is also evident at the transcriptomic scale, with greater enrichment of differentially expressed genes in the presence of multiple tumor-mimicking ECM cues.

However, MCF-10A cells did not exhibit a similar transcriptional response, underscoring the cell-line specific differences in how multiple mechanical signals are integrated during invasion in complex microenvironments. Future studies investigating which ECM cues most critically drive invasive behavior across different cell lines could provide valuable insights for developing targeted therapeutic strategies in breast cancer treatment.

During cancer invasion and metastasis, the migration of tumor cells has been shown to be regulated by various physical cues within the tumor microenvironment^10,74,75^. In this study, we observed that matrix stiffness, stress relaxation, and collagen density together dictate cell migration. For instance, in soft matrices, MDA-MB-231 cells migrate with greater speed in the presence of fast stress relaxation, however, in stiff matrices, this trend is reversed.

These observations align with previous findings reporting enhanced migration of MCF-10A cells on soft, fibronectin-coated viscoelastic substrates but reduced migration on stiff substrates^76^. However, a recent report showed that MDA-MB-231 cells on Matrigel-coated 2D substrates demonstrated increased migration in response to faster stress relaxation on substrates with a Young’s modulus similar to our stiff matrices^17^. These differences might be attributed to differing modes of migration in 2D versus 3D and laminin/collagen IV ligands employed in their study, known to trigger distinct integrin-mediated signaling pathways^77^. Such variations underscore the importance of matrix composition, dimensionality, and ligand specificity in modulating cell migration behavior.

It is well established that breast tumors are stiffer than healthy breast tissue, and this increased stiffness is associated with the presence of malignant traits in cells and poor patient survival^78,79^. Recent studies have also shown that breast cancer progression is linked to changes in tissue viscoelasticity^9–11^. Although extensive research has focused on elucidating the pathways involved in mechanotransduction in response to varying elastic moduli during disease progression, how cells sense and respond to changes in ECM viscoelasticity remains less well understood. To the best of our knowledge, our study is the first to demonstrate that Sp1 signaling plays a critical role in regulating the cellular response to matrix stress relaxation. Further, our observation that Sp1 target genes are upregulated across fast relaxing conditions (with the exception of Soft-Fast-Col_low_ condition) and are associated with poor patient survival, underscores the potential role of Sp1 mechanotransduction in driving disease progression.

High expression of Sp1 is known to correlate with poor prognosis and shorter survival times in multiple cancers, including breast cancer^57,80^. Further, Sp1 undergoes a large number of post-translational modifications that regulate its stability and transcriptional activity^57^.

Previous work has shown that Sp1-HDAC3/8 signaling can drive the malignant phenotype via changes in chromatin accessibility in response to increased ECM stiffness in mammary epithelial cells^16^. Another study has previously reported that direct force application to fibroblasts via collagen-coated beads led to higher levels of phospho-Sp1^58^. However, the role of Sp1 and its downstream modifications in response to matrix mechanical remodeling during cancer progression remains understudied. ERK1/2 acts as a downstream effector in Ras signaling that can shuttle into the nucleus^81,82^ and phosphorylate multiple transcription factors such as Sp1 and HIF1A, c-Myc, STATs, Jun, and Fos^64,65,83^. PI3K also has been shown to phosphorylate Sp1 and regulate downstream gene expression^67,84^. We demonstrate that Sp1 phosphorylation through ERK1/2 and PI3K signaling pathways regulates the cell invasion in response to matrix mechanics. Our findings highlight the importance of studying Sp1 signaling as a novel candidate toward developing therapies targeting cell response in response to ECM remodeling.

We also observed loss of the invasive morphologies, as well as downregulation in nuclear localization of phospho-Sp1 in both MDA-MB-231 and MCF-10A cells upon inhibiting actin-polymerization in stiff matrices. It is known that ERK can interact with the actomyosin cytoskeleton and favor actin polymerization^85^. Chemical and optogenetic induction of cell protrusions, which are characterized by increased actin polymerization, have also been shown to activate ERK signaling^86^, while inhibition of actomyosin contractility can abrogate downstream ERK signaling^87^. These findings might also help explain changes in nuclear localization of phospho-Sp1. Intriguingly, in MDA-MB-231 cells, we did not observe changes in phospho-Sp1 nuclear localization in Soft-Fast matrices. Future work is needed to determine the role of other cytoskeletal elements, or their crosstalk in regulating Sp1 signaling in these matrices, to better understand how ECM mechanics influence tumor progression and therapeutic responses.

In summary, our work demonstrates that various ECM cues within the complex tumor microenvironment can concurrently influence cells to drive diverse malignant traits. We identified Sp1 and its phosphorylation via ERK and PI3K signaling as a novel regulator of invasion in response to multiple tumor-mimicking mechanical cues, as well as its dependence on the cytoskeletal network. Furthermore, we also identified Sp1 target genes enriched across fast stress relaxing conditions, whose upregulation correlated with poor survival for basal-subtype patients. Our work thus highlights the need to further investigate the mechanism of Sp1 mechanosignaling for therapeutic applications.

## Methods

### Preparing Alginate-collagen matrices

PronovaⓇ UP-VLVG (MW < 75 kDa) and LF20/40 (265 kDa) sodium alginates were used for producing fast and slow relaxing matrices, respectively. Calcium sulfate was used to ionically crosslink the alginate chains. 0.5 mg/ml or 2 mg/ml rat tail collagen I (Advanced Biomatrix) was incorporated to form the alginate-collagen interpenetrating network (IPN) matrices. Briefly, the alginate solution in a Luer-lock syringe was mixed with collagen I and calcium sulfate solutions diluted in DMEM or DMEM/F12 in a second Luer-lock syringe.

Collagen was neutralized using 1 M NaOH prior to mixing with the alginate solution. The syringes were then coupled using a Luer-lock and the solution was mixed back and forth 20 times before depositing it into well plates. The solution was allowed to gel for 2 hrs at 37°C. The exact recipe for each matrix condition can be found in Supp. Table 1.

### Single cell encapsulation and cell culture

MCF-7 and MDA-MB-231 cells were obtained from ATCCⓇ and cultured in media with DMEM (4.5 g/L D-glucose, ThermoFisher Scientific), 10% fetal bovine serum (ThermoFisher Scientific), and 1% Penicillin/Streptomycin (ThermoFisher Scientific). Cells were used up to passage 40. Cells were encapsulated at 250,000 cells/ml and cultured in gels for 7 days.

MCF-10A cells were obtained from ATCCⓇ and cultured in media containing DMEM/F12 (ThermoFisher Scientific), 5% horse serum (ThermoFisher Scientific), 1% Penicillin/Streptomycin (ThermoFisher Scientific), 20 ng/ml epidermal growth factor (ThermoFisher Scientific), 0.5 mg/ml hydrocortisone (Sigma Aldrich), 100 ng/ml cholera toxin (Sigma Aldrich), and 10 μg/ml insulin (Sigma Aldrich). MCF-10A cells were encapsulated at a density of 100,000 cells/ml. Media for all cell types was replenished every 2-3 days.

### Mechanical characterization of hydrogel matrices

Gels were characterized using an Anton-Paar MCR-502e strain-controlled rheometer. The gel volume was cast between 25 mm diameter plates and a gap of ∼1.2 mm. Mineral oil was applied to the edges of the plates to minimize solvent evaporation. The storage modulus was measured using a 0.5% strain and a frequency of 1 Hz until it reached a steady-state value. For measuring stress relaxation, a 10% strain was applied, and the time required to reach 50% of the maximum stress was recorded.

### Spheroid invasion assay

Spheroids were formed using the hanging-drop method. Collagen I was added to cell suspensions at a concentration of 6.75 μg/ml. 10 μl droplets containing 3,000 cells/drop were deposited on petri dish lids. The lids were then inverted, and the dishes were filled with 1X PBS. Spheroids were allowed to form for 24-36 hrs. Each spheroid was then collected manually and encapsulated into alginate-collagen matrices. MDA-MB-231 spheroids were cultured in gels for 3 days, while MCF-10A spheroids were cultured for 7 days.

### Murine mammary organoid culture

Mammary organoids were extracted in accordance with a previous protocol published^42^. Briefly, mammary glands from 10-week-old female C57BL/6 strain mice were minced and digested in a collagenase solution on a shaker. 10 ml collagenase solution was prepared by combining 9 ml DMEM/F12 (Fisher #11320082), 0.5 ml fetal bovine serum (Gibco), 5 μl insulin (Sigma, Cat#I-1882), 10 μl gentamicin (50 mg/ml stock, ThermoFisher Cat#15750-060), 200 μl collagenase (100 mg/ml stock, Sigma #2139), and 200 μl trypsin (100 mg/ml stock, Sigma #T7409). The solution was then centrifuged, and the fatty layer was transferred to a BSA-coated tube. The fatty layer was resuspended in DMEM/F12, centrifuged, and the supernatant was discarded. This pellet was combined with the original pellet in a tube and resuspended in 4 ml of DMEM/F12. 40 μl DNase (Sigma Cat#D4263) was added to this suspension, centrifuged, and the supernatant was discarded. The pellet was washed 4X with DMEM/F12 by pulsing it for 3-4s in a centrifuge. Post extraction, organoids were encapsulated in alginate-collagen matrices and maintained in a DMEM/F12 media containing 1% penicillin/streptomycin, 1% insulin-transferin-selenium-X (ITS)(Gibco), and 2.5 nM FGF2 (Sigma).

### Cell migration assay

MDA-MB-231 cells were stained with 1 μg/ml R18 cell membrane dye (ThermoFisher Scientific) for 1 hr before encapsulation into alginate-collagen matrices. Gels containing cells were cast into 8 well glass-bottom live-cell chambers (Labtek). Sterile agarose pieces were used to prevent gels from floating. Live-cell microscopy (Leica SP8 confocal microscope) was performed on day 2 post-encapsulation using a 10x air objective. Cell migration within gels was recorded over a 100 μm thick stack for ∼16 hrs. Maximum Z-projection was performed and then cell migration average speed and mean-squared displacement were measured using Imaris software. A custom MATLAB script was used to plot individual cell 2D trajectories.

### RNA-sequencing and analysis

Cells were extracted from alginate-collagen matrices by rocking them in Falcon tubes containing 2.5 mg/ml collagenase (Sigma, Cat#C0130) solution in PBS for 30 mins at 37°C. After this, the tubes were centrifuged, and the supernatant was removed. Following this, the pellet was dissolved in 10 ml ice-cold EDTA (50 mM) and placed on a rotator for 10 mins.

The tubes were centrifuged, and the supernatant was then removed. The cell pellet was then lysed using Trizol (Life Technologies) and RNA was extracted using the total RNA mini prep kit according to the manufacturer’s instructions (Epoch Life Sciences).

Bulk-mRNA-sequencing library prep was done using Cel-Seq2 pipeline as described previously^88,89^. Briefly, 10 ng RNA was reverse-transcribed using the CelSeq2 RT-primer, DTT (0.1 M), dNTPs (New England Biolabs, Cat# N0447l), and Superscript II reverse transcriptase (Invitrogen, Cat# 18064014), followed by second-strand synthesis using RNAseH (ThermoFisher Scientific, Cat# EN0202), E.Coli DNA Pol I (Invitrogen, Cat# 18010025), E.Coli ligase (Invitrogen, Cat#18052-019), and the second strand buffer (Invitrogen, #18052-019). After cDNA cleanup using DNA beads (AMPure, Cat# A63882), in vitro transcription was performed using the MEGAscript T7 kit (Invitrogen, Cat# A57622) resulting in amplified single stranded RNA (aRNA). After aRNA treatment with ExoSAP-IT PCR reagent (ThermoFisher, 78200), RNA strands were fragmented using a fragmentation buffer containing 200 mM Tris-acetate (pH 8.1), 500 mM KOAc and 150 mM MgOAc. aRNA cleanup was then performed using RNA-beads (AMPure, Cat#A63987). Next, aRNA was reverse transcribed using a RT random hex primer and cDNA was then amplified using a RNA PCR primer (RPI), a uniquely indexed Illumina primer, and a PCR master mix (New England, Cat#M0541S). After bead cleanup and quality control using a bioanalyzer (Agilent), DNA libraries were sequenced using NovaSeq^TM^ 6000 system (Illumina). The raw reads were then mapped to the hg19 (GRCh37) reference genome.

Differentially expressed genes were determined using the DESeq2 library in R. A minimum log-fold change value of 1 and an adjusted p-value cutoff value of 0.05 were used in the analysis. Volcano plots were made using the ggplot package in R. Gene Set Enrichment Analysis (GSEA) was performed to determine differentially enriched oncogenic pathways within the C6: oncogenic gene sets in GSEA software. Transcription factor enrichment analysis was performed using the TRRUST database. Sp1 target genes were chosen from TRRUST database. Transcription factors with adjusted p-values < 0.05 were considered. Principal component analysis (PCA) was performed using the PlotPCA function in the DEseq library. Sp1 target gene heatmap and clustering was performed using the pheatmap library in R. Sp1 genes implicated in breast neoplasm and neoplasm metastasis were obtained from the TRRUST database. Kaplan-Meier plots were made using the survival and survminer libraries in R.

### Immunofluorescence imaging

Cell culture media was removed and replaced with 4% paraformaldehyde. After fixing for 45 mins at 37°C, gels were washed 2-3X with DPBS containing Ca^2+^/Mg^2+^. Gels were then dehydrated overnight at 4°C in a 30% sucrose solution made in DPBS containing Ca^2+^/Mg^2+^. The next day, gels were incubated in a 1:1 solution of OCT (Fisher Scientific) and 30% sucrose solution for 4-6 hrs at 4°C. The gels were then frozen in cryomolds in OCT over dry ice. 40 μm thick gel sections were cut using a cryostat (Leica CM1850) and placed on poly-L-lysine coated glass slides.

Sections were washed in DPBS containing Ca^2+^/Mg^2+^ for 1h at room temperature and then blocked for 1 hr at room temperature using a solution of 1X DPBS (with Ca^2+^/Mg^2+^), 1% bovine serum albumin, 0.1% Triton X-100, 0.3 M glycine, 10% goat serum, and 0.05% sodium azide. Next, sections were incubated with primary antibody solutions overnight at 4°C. Sections were then washed 3X using a blocking buffer and incubated in Alexa Fluor 488 antibody solution (1:1000 dilution), DAPI (1 μg/ml) and Alexa-Fluor 647 Phalloidin (1:100, ThermoFisher Scientific). Next, slides were washed 3X with the blocking buffer, mounted using Prolong Gold antifade reagent (ThermoFisher Scientific) and their edges were sealed using nail polish. Antibodies used were anti-phospho-Sp1(T453) (Abcam ab59257) and Alexa Fluor 488 goat anti-rabbit IgG (ThermoFisher, Cat#A11998). Samples were imaged using a Leica SP8 confocal microscope.

### Collagen fiber alignment

Samples were imaged using the 25X objective on a Leica SP8 confocal microscope using the 488 nm laser in the reflectance mode. Fiber dimensions (length and width), as well as fiber orientation relative to cell boundaries was analyzed using CT-Fire, a freely available software written in MATLAB script^90^. Fibers selected were within 200 μm from the cell boundary. Fibers oriented at an angle greater than 70° with respect to the cell boundary were categorized as aligned fibers.

### Pharmacological Inhibition

Small molecule inhibitors were dissolved in dimethyl sulfoxide (DMSO) and diluted in cell culture media. The inhibitors used were ML-7 (25 μM, Cayman Chemical, #11801), Cytochalasin-D (1 μM, Cayman Chemical, #11330), LY294002 (20 μM, Cayman Chemical, #70920), SCH772984 (0.1 μM, Cayman Chemical, #19166), Mithramycin A (50 nM, Cayman Chemical, #11434). A DMSO vehicle was used for the control samples.

### Image analysis

Cells stained with phalloidin were used to quantify roundness. Images were thresholded and then analyzed using ImageJ particle analysis feature. For spheroid invasion assays, spheroids were stained with R18 membrane dye and their invasion boundaries within the matrix were manually traced. The area and circularity of these invasion boundaries were then quantified using ImageJ. For analyzing the nuclear localization of phospho-Sp1, a nuclear mask using the DAPI stain was first created using a custom MATLAB code.

Multiplying the phospho-Sp1 image channel with the nuclear mask resulted in the phospho-Sp1 within the nucleus. Similarly, a cytoplasmic mask was created using the phalloidin stain and then subtracting the nuclear mask from it. Cytoplasmic phospho-Sp1 was similarly obtained by multiplying the binary cytoplasm mask with the phospho-Sp1 image. Mean intensity of the nuclear and cytoplasmic phospho-Sp1 images was then calculated using ImageJ. Ratio of the mean nuclear and cytoplasmic intensity of phospho-Sp1 for each image was used to quantify the nuclear localization.

### Statistical Analysis

All statistical analyses were done using GraphPad (Prism) v.10.2. Specific tests used for analysis can be found in the figure descriptions.

## Acknowledgments

We thank members of the Stowers lab for critical feedback on this manuscript. We thank Megan D. Li and Sabrina M. Jackson for help with sample preparation for confocal imaging. We thank Alban Sauret Lab at UCSB for access to the rheometer. We acknowledge Meghan Morrisey lab at UCSB for access to mouse mammary tissue. This work was supported by a UC Cancer Research Coordinating Committee grant to R.S.S. (C22CR4078), a Breast Cancer Alliance Young Investigator Grant to R.S.S. and an American Cancer Society Catalyst award to R.S.S. (CAT-24-1379187-01-CAT). A.S. was supported by the California Institute of Regenerative Medicine (CIRM) EDUC4 training grant. S.S.D. was supported by NIH grants R01HD099517 and R01HG011013, and an NSF CAREER grant (Award # 2339849). Schematic in Fig. 1A was created with <u>BioRender.com</u>. We acknowledge the use of the NRI-MCDB Microscopy Facility at University of California, Santa Barbara.

## Author contributions

Conceptualization, A.S., R.S.S.,S.S.D.; methodology, A.S., R.S.S., S.S.D.; software, A.S., R.F.S., K.A.H.; validation, A.S., R.F.S., J.M.L., J.A.B.; formal analysis, A.S., R.F.S., K.A.H.; investigation, A.S., R.F.S., J.M.L., J.A.B., K.A.H.; resources, R.S.S.; writing – original draft, A.S., R.S.S.; writing – review and editing, A.S., R.S.S., S.S.D., R.F.S., J.M.L., J.A.B., K.A.H.; visualization, A.S., J.A.B., R.S.S.; supervision, R.S.S.; funding acquisition, R.S.S.

## Competing Interests

The authors declare no competing interests.

## Notes

### Competing Interest Statement

The authors have declared no competing interest.

