## Supplemental Information for "Sp1 mechanotransduction regulates breast cancer cell invasion in response to multiple tumor-mimicking extracellular matrix cues"

**Supplementary Table 1:** Alginate-collagen gel recipes for soft/stiff, slow/fast relaxing, and $Col_{low}$/$Col_{high}$ conditions

| **Matrix condition** | **Sodium alginate** | **Collagen I** | **$CaSO_4$** |
|---|---|---|---|
| Soft-Slow-$Col_{low}$ | 5 mg/ml LF20/40 | 0.5 mg/ml | 2 mM |
| Stiff-Slow-$Col_{low}$ | 5 mg/ml LF20/40 | 0.5 mg/ml | 21 mM |
| Soft-Fast-$Col_{low}$ | 10 mg/ml UPVLVG | 0.5 mg/ml | 5 mM |
| Stiff-Fast-$Col_{low}$ | 10 mg/ml UPVLVG | 0.5 mg/ml | 17 mM |
| Soft-Slow-$Col_{high}$ | 5 mg/ml LF20/40 | 2 mg/ml | 2 mM |
| Stiff-Slow-$Col_{high}$ | 5 mg/ml LF20/40 | 2 mg/ml | 21 mM |
| Soft-Fast-$Col_{high}$ | 10 mg/ml UPVLVG | 2 mg/ml | 2 mM |
| Stiff-Fast-$Col_{high}$ | 10 mg/ml UPVLVG | 2 mg/ml | 10 mM |

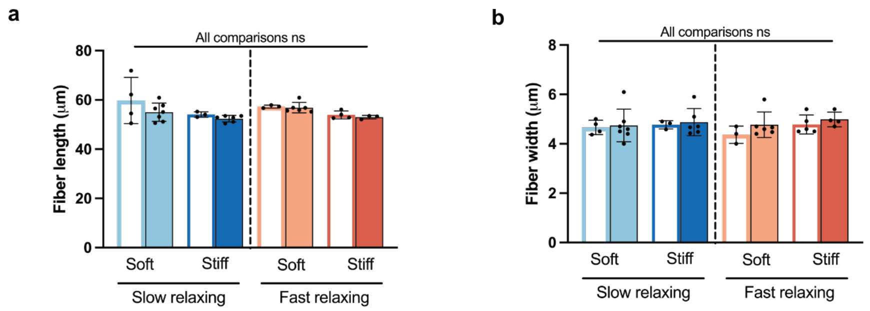


**Supplementary Fig. 1: Collagen fiber dimensions are not significantly different across matrices. a, b** Quantification of collagen fiber length and width across all 8 matrix conditions shows no significant differences (n ≥ 3; mean ± s.d.; ANOVA with Tukey's multiple comparison test). not significant = ns.

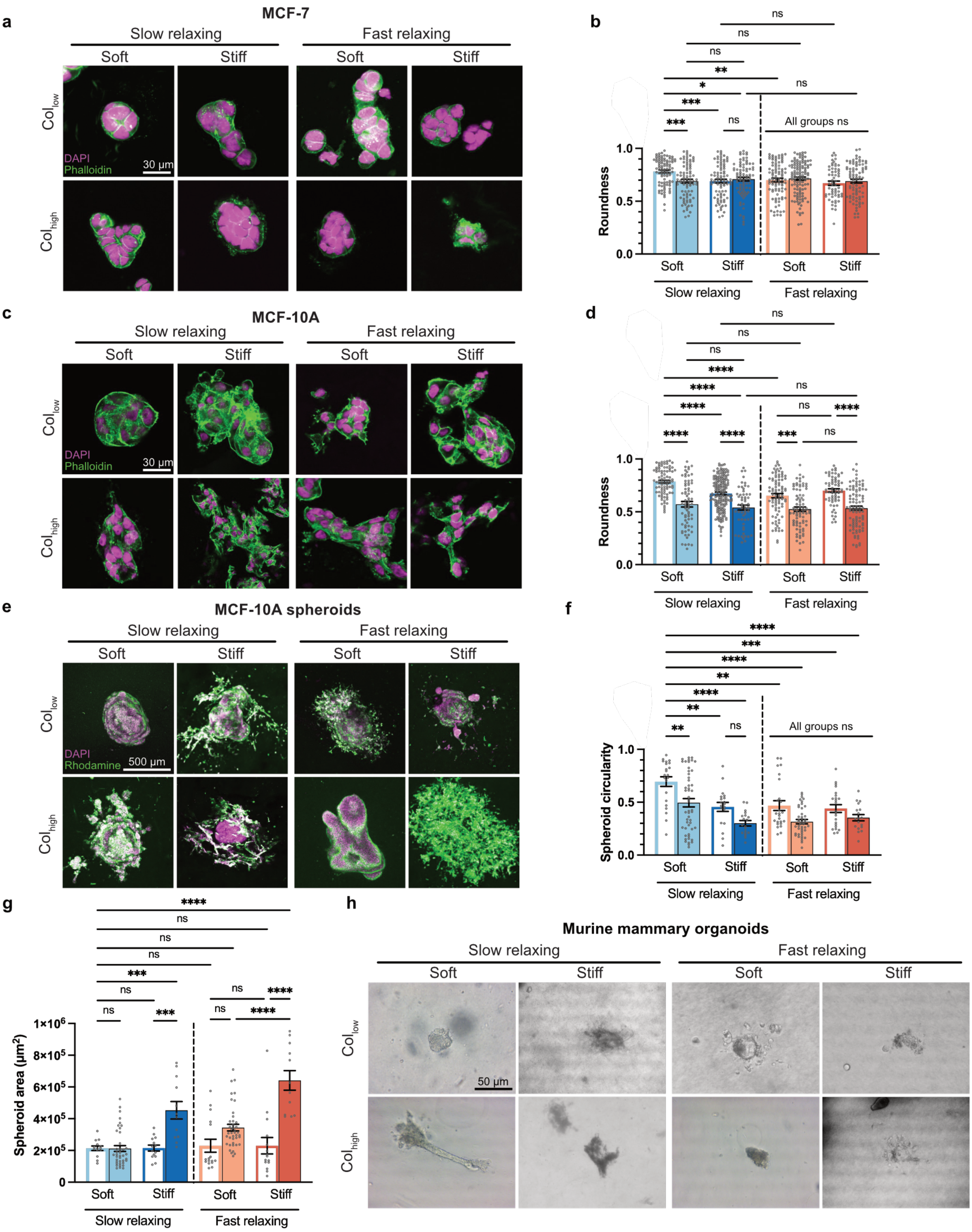


**Supplementary Fig. 2: MCF-7, MCF-10A, and mouse mammary organoids show diverse morphologies in the presence of individual and multiple tumor-mimicking ECM cues. a,** Immunofluorescence imaging of phalloidin-stained MCF-7 cells and **b,** quantification of cell cluster roundness. **c,** Immunofluorescence imaging of phalloidin-stained MCF-10A cells and **d,** quantification of cell cluster roundness. Individual or multiple cues led to significant decrease in roundness. **e,** Rhodamine-stained MCF-10A spheroids and **f,** quantification of spheroid circularity and **g,** spheroid area. The presence of individual or multiple cues also led to significant reduction in circularity. **h,** Bright-field imaging of murine mammary organoids encapsulated showed that the organoids remained intact only in Soft-Slow-$Col_{low}$ condition. For **b, d,** n ≥ 20 cells per replicate, with 3 independent replicates; mean ± s.e.m.; Welch ANOVA test with Dunnett's multiple comparison test. For **f, g,** n ≥ 5 spheroids per replicate, with 3 independent replicates; mean ± s.e.m., ANOVA test with Tukey's multiple comparison test. *p < 0.05; **p < 0.01; ***p < 0.001; ****p < 0.0001, not significant = ns.

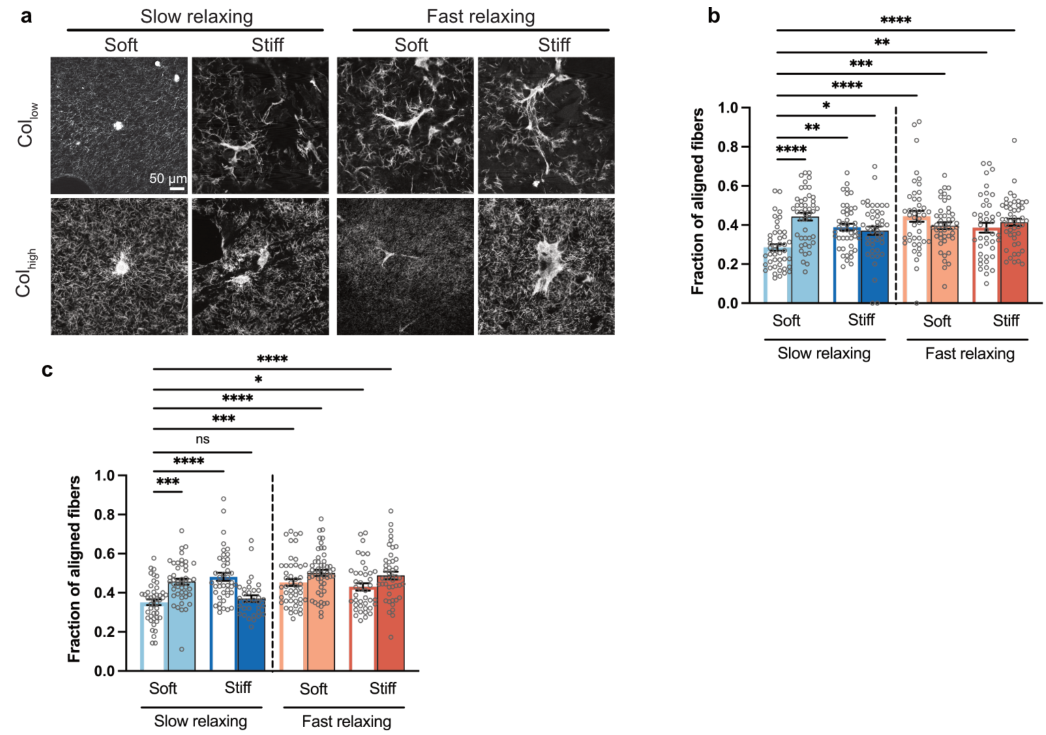


**Supplementary Fig. 3: Cells exhibit increased collagen fiber alignment in presence of tumor-mimicking ECM cues. a,** Confocal reflectance micrographs of MDA-MB-231 cells encapsulated in alginate-collagen matrices. **b,** Alignment of collagen fibers with respect to MDA-MB-231 cell boundary in regions of interest within 200 μm from the cell boundary. The percentage of fibers aligned at greater than 70° was quantified. Compared to Soft-Slow-$Col_{low}$ condition, all other conditions showed a significantly higher proportion of aligned collagen fibers. **c,** MCF-10A cells also showed a significantly higher fraction of aligned collagen fibers, except in the Stiff-Fast-$Col_{low}$ condition. For both **b, c,** n ≥ 10 cells per replicate, with 3 independent replicates; mean ± s.e.m., ANOVA test with Dunnett's multiple comparison test. *$p < 0.05$; **$p < 0.01$; ***$p < 0.001$; ****$p < 0.0001$, ns = not significant.

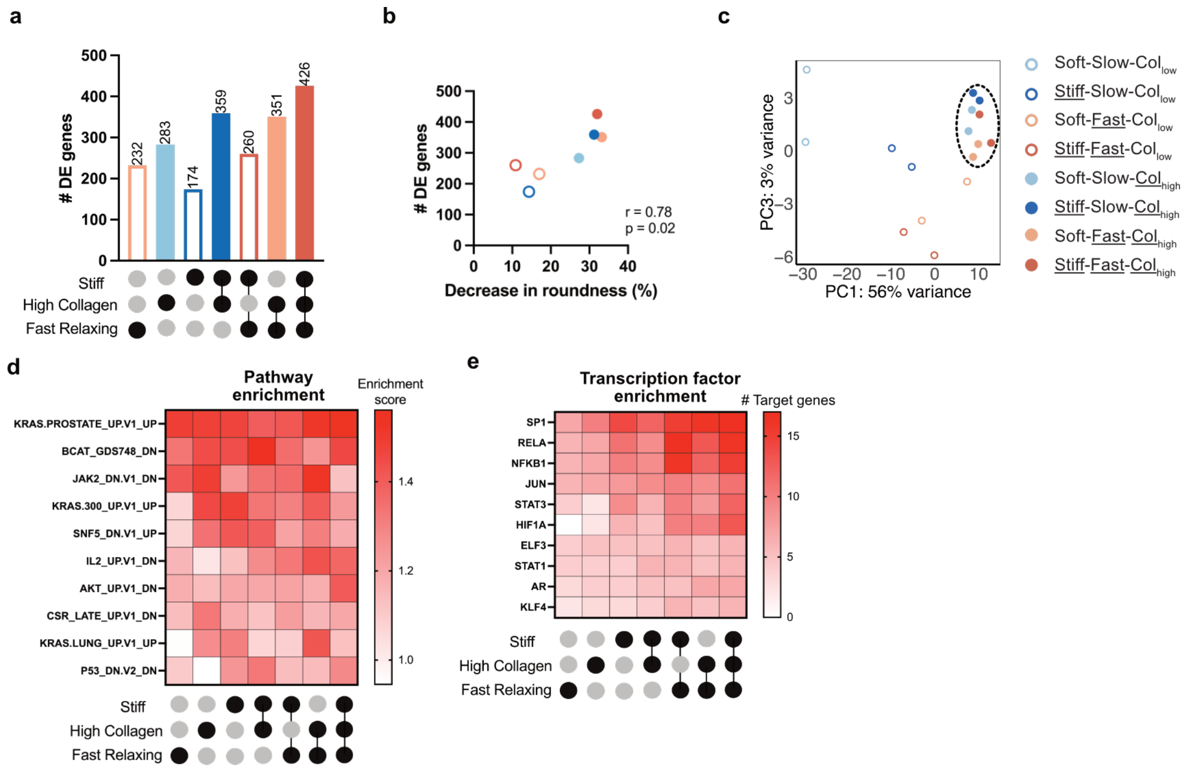


**Supplementary Fig. 4: The presence of invasive matrix cues is associated with enrichment of Sp1 target genes also in MCF-10A cells. a,** The number of differentially expressed genes enriched in MCF-10A cells in various matrix conditions with respect to Soft-Slow-Col$_{low}$ condition (n = 2 replicates). **b,** Plots showing a positive correlation between the decrease in cluster roundness and the number of differentially expressed genes. The p value and correlation coefficient were calculated using one-tailed Spearman correlation test. **c,** PCA plot shows a clustering of all high collagen conditions. **d,** GSEA oncogenic pathway enrichment analysis of differentially expressed genes shows enrichment of KRAS signaling. **e,** TRRUST analysis of differentially expressed genes shows enrichment of Sp1 target genes in all invasion-inducing matrix conditions.

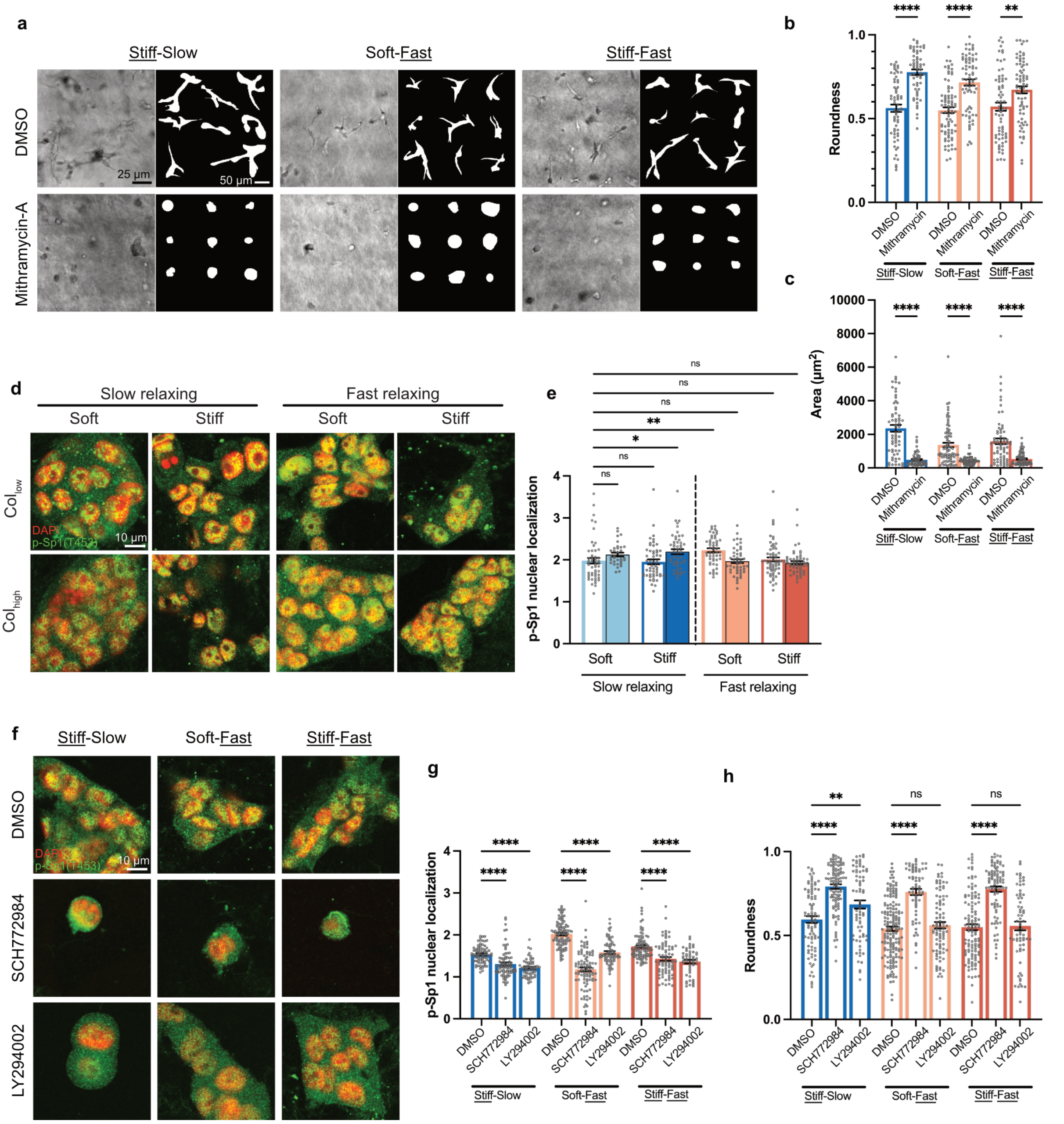


**Supplementary Fig. 5: Sp1 regulates the invasive phenotype in response to multiple tumor-mimicking ECM cues in MCF-10A cells. a,** Inhibiting Sp1 activity using mithramycin-A reverts the invasive morphologies to round clusters in Stiff-Slow, Soft-Fast, as well as Stiff-Fast matrix conditions. **b, c,** Quantification of the change in roundness and cluster area upon Sp1 inhibition. A significant increase in roundness and decrease in area is seen in all 3 conditions (n ≥ 15 cells per replicate, with 3 independent replicates; ANOVA with Sidak's multiple comparison test). **d,** Confocal immunofluorescence imaging and **e,** Quantification of p-Sp1(T453) nuclear localization did not show a significant difference in most conditions (n ≥ 15 cells per replicate, with 3 independent replicates; mean ± s.e.m; ANOVA with Dunnett's multiple comparison test). **f,** Sp1 phosphorylation upon inhibition using SCH772984 (ERK1/2) and LY294002 (PI3K) in Stiff-Slow, Soft-Fast, as well as Stiff-Fast matrix conditions. **g,** Both ERK1/2 and PI3K inhibition led to a significant decrease in nuclear localization of p-Sp1(T453). **h,** While ERK1/2 inhibition led to significant decrease in roundness in all conditions, PI3K inhibition had a significant effect only in the Stiff-Slow matrix condition. For both **g, h,** n ≥ 15 cells per replicate, with 3 independent replicates; mean ± s.e.m.; ANOVA with Sidak's multiple comparison test. *p < 0.05; **p < 0.01; ***p < 0.001; ****p < 0.0001, not significant = ns. For **a, b, c, f, g, h,** all 3 conditions have a low collagen density.

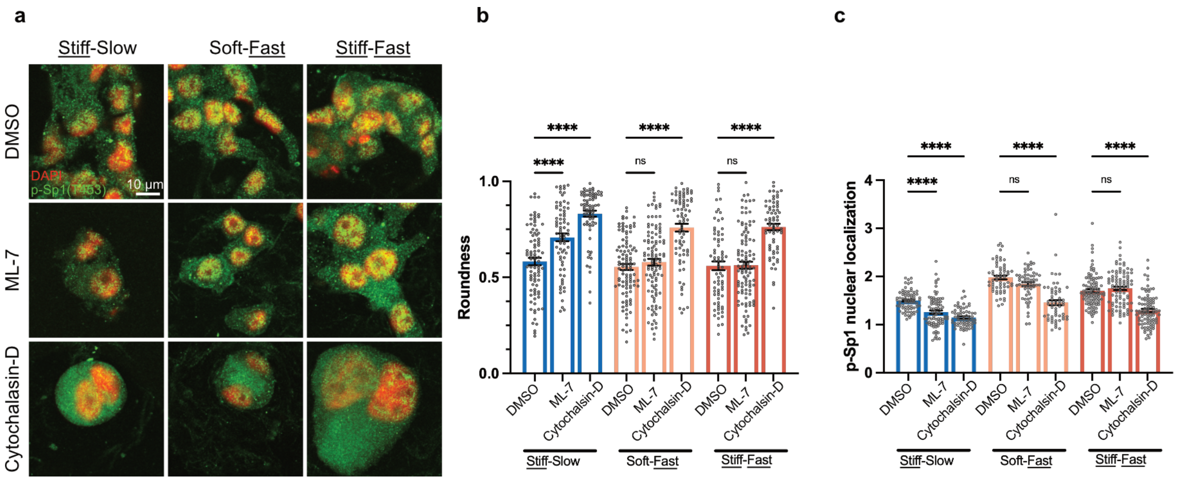


**Supplementary Fig. 6: Actin polymerization regulates p-Sp1 nuclear localization in MCF-10A cells. a,** Myosin light chain kinase (MLCK) inhibition via ML-7 and actin polymerization inhibition via cytochalasin-D in MCF-10A cells encapsulated in Stiff-Slow, Soft-Fast, as well as Stiff-Fast matrix conditions. Immunofluorescence imaging was performed on DAPI and p-Sp1(T453) stained cells. **b,** MLCK inhibition led to a significant increase in cluster roundness only in the slow relaxing condition but inhibiting actin polymerization led to significant increase in roundness in all 3 conditions. **c,** Actin polymerization inhibition led to a significant decrease in nuclear localization levels of p-Sp1 in all conditions. For **b, c,** $n \geq 15$ cells per replicate, with 3 independent replicates; mean ± s.e.m.; ANOVA with Sidak's multiple comparison test. *$p < 0.05$; **$p < 0.01$; ***$p < 0.001$; ****$p < 0.0001$, not significant = ns.
